# Partial Inhibition of Viral Replication Machinery Enhances Recombination in Herpes Simplex Viruses

**DOI:** 10.1101/2025.09.15.676234

**Authors:** Maya Ralph-Altman, Daniel Avhar, Yuval Altman, Itai Cohen, Hadas Azulay, Tal Korner, Sireen Sweed, Enosh Tomer, Oren Kobiler

## Abstract

Herpes simplex viruses (HSV-1 and HSV-2) are widespread human pathogens, most commonly causing oral and genital lesions. These DNA viruses use recombination as a major driver of evolution. Intragenomic and intergenomic recombination events can be detected both in vivo and in vitro. As viral recombination is tightly linked to replication, deciphering mechanisms that specifically effect recombination remains a challenge. Here, we employed a triple-fluorescent color recombination assay to identify homology mediated and non-canonical recombination events between co-infecting HSV-1 strains. We developed a deep learning model that detects and classifies progeny plaques according to their colors. This setup enabled us to perturb the infection process using small molecule inhibitors targeting either viral or host proteins. We identified that inhibitors that reduce infectious viral progeny, increased viral homology mediated recombination. The antiviral drugs, including acyclovir, also increased recombination between HSV-1 and HSV-2 and aberrations in the progeny viral genomes. Taken together our results indicate that commonly used anti-herpes drugs increase intraspecies and interspecies recombination rates and genetic rearrangements.

**Significance:** Herpes simplex viruses cause significant morbidity in all human populations. Current therapies rely primarily on antivirals that target viral replication. Viral recombination plays a key role in replication and evolution of these dsDNA viruses. To better characterize the viral recombination process, we introduce a fluorescence-based HSV-1 recombination assay coupled with deep learning-based plaque classification, enabling high-throughput quantification of recombination rates. We found that clinically relevant moderate concentrations of commonly used antiviral replication inhibitors, enhance homology-mediated intraspecies recombination, HSV-1/HSV-2 interspecies recombination, as well as the accumulation of defective genomes. We conclude that partial inhibition of the herpes replication complex can promote viral diversification, with potential implications for HSV evolution and drug resistance.

## Introduction

HSV-1 and HSV-2 are ubiquitous pathogens, estimated to infect 60% and 30% of the human population, respectively (1). While most infections are asymptomatic, many people suffer from common herpetic lesions (cold sores) in the facial or genital areas. Herpes simplex viruses (HSV) are probably the most common sexually transmitted disease worldwide (2). It was originally thought that lesions caused by HSV-1 and HSV-2 occurred primarily in the facial and genital areas, respectively. However, recent studies have found that HSV-1 can be detected in approximately 30% of genital herpes cases (1, 3). In newborns, immunocompromised people, and rarely in otherwise healthy individuals, HSV can cause severe encephalitis and other life-threatening diseases. Currently, no vaccines are approved for use, and only few antiviral drugs are in clinical use (4, 5).

HSV-1 and HSV-2 have distinct evolutionary histories (6), yet the genetic organization and most viral-encoded proteins are homologous. Amino acid identity between orthologous proteins ranges from ∼45% to 90% (7), with the viral DNA replication machinery being highly conserved, and the exposed structural glycoproteins showing the greatest divergence. Viruses enhance the probability of adapting to changing environments by maintaining genetic diversity. Genetic diversification can arise from point mutations or from recombination events and can lead to more virulent or drug-resistant strains. Sequencing analyses of clinical isolates of HSV indicate frequent recombination; and notably, even recombination events between HSV-1 and HSV-2 have been detected (7, 8), supporting the view that recombination is a major evolutionary driver herpesvirus diversity (9, 10). Both intergenomic and intragenomic recombination events are common during viral replication in vitro and can be observed in clinical samples (10, 11).

The herpes simplex genomes contain two unique regions (U_L_ and U_S_), each flanked by repeat regions. Three copies of the *a* sequence are found at the end of the genomes and at the junction of the internal repeats. We have recently characterized the sites in which non-canonical rearrangements occur at the HSV-1 genomes (11). We identified that many of these rearrangements occur within or adjacent to the repeat regions of viral genomes. Short repeats or short homology sequences were enriched at the sites of these rearrangements; however, rearrangements could be detected without any detectable homology (11).

Like all herpesviruses, HSV replicates its large double-strand DNA genome in distinct replication compartments (RCs) within the host nucleus (12). In the RC’s, both RNA transcription and DNA replication take place. Viral DNA replication is carried out by the viral replication machinery, including the viral encoded polymerase and helicase (13), which are the targets of currently available drugs (5). RCs usually originate from a single incoming viral genome, and only limited number of them can be detected per cell (14, 15). As RC’s are dynamic structures, they grow, move and coalesce during the replication process (16). We have shown that collision between RCs provide the opportunity for intergenomic recombination to take place (17).

Both viral and host proteins are recruited to the viral RCs to allow efficient viral replication (13, 18). Most Viral proteins involved in replication have been identified. Two viral proteins: the single-strand DNA binding protein, ICP8 and the alkaline nuclease, UL12 form a complex that promote recombination via the single-stand annealing mechanism (19, 20). It was suggested that viral replication is enhanced by recombination dependent replication in which branching concatemers are generated via recombination processes (21–23).

Recent studies have identified several transcription, replication and restriction host factors associated with viral replication forks (24–26). Some of these proteins are essential for efficient viral replication, while others are thought to contribute to host antiviral processes by repressing viral transcription and replication (27). However, the precise role of most of these proteins during viral replication remains unclear. Additionally, some of these proteins are thought to play a role in viral recombination processes (28).

The central role of recombination in herpesvirus biology prompted the development of fluorescent proteins–based assay capable of detecting both intergenomic homology-mediated and non-canonical recombination events. Using this system, we show that inhibition of the viral replication machinery by moderate concentrations of commonly used antiviral drugs increases recombination rates while reducing viral infectivity.

## Results

### Construction of HSV-1 fluorescence-based recombination system

To identify specific mechanisms involved in HSV genomes recombination, we developed an assay (**Figure 1A**) for detection and quantification of homology-mediated recombination (HMR) and non-canonical recombination (NCR) events that include deletions, duplications or other rearrangements. The assay is based on two derivatives of HSV-1 strain 17: HSV-1 OK11 (29), expressing the mCherry (RFP) gene located between UL37 and UL38 viral genes, and HSV-1 OK42 (30) expressing both mTurq2 (CFP) and mVenus (YFP) genes located between UL37 and UL38 genes and fused within the UL25 gene, respectively (**Figure 1B**). Coinfection with these two viral derivatives expressing fluorescent proteins (FPs) produced seven distinct progeny types, each characterized by a unique FP pattern: two parental viruses, two recombinants likely arising from intergenomic HMR, and three recombinants most consistent with intergenomic NCR events (**Figure 1C**). The single-color mTurq2 plaques, that we classify as a HMR event, may theoretically also result from intragenomic deletion. However, as the single-color mVenus plaques were the least frequent across all experiments (**Figure 1D**) and while setting up the assay, we did not detect any colorless plaques, we suggest that intragenomic deletions are a relatively rare event.

**Figure 1.**
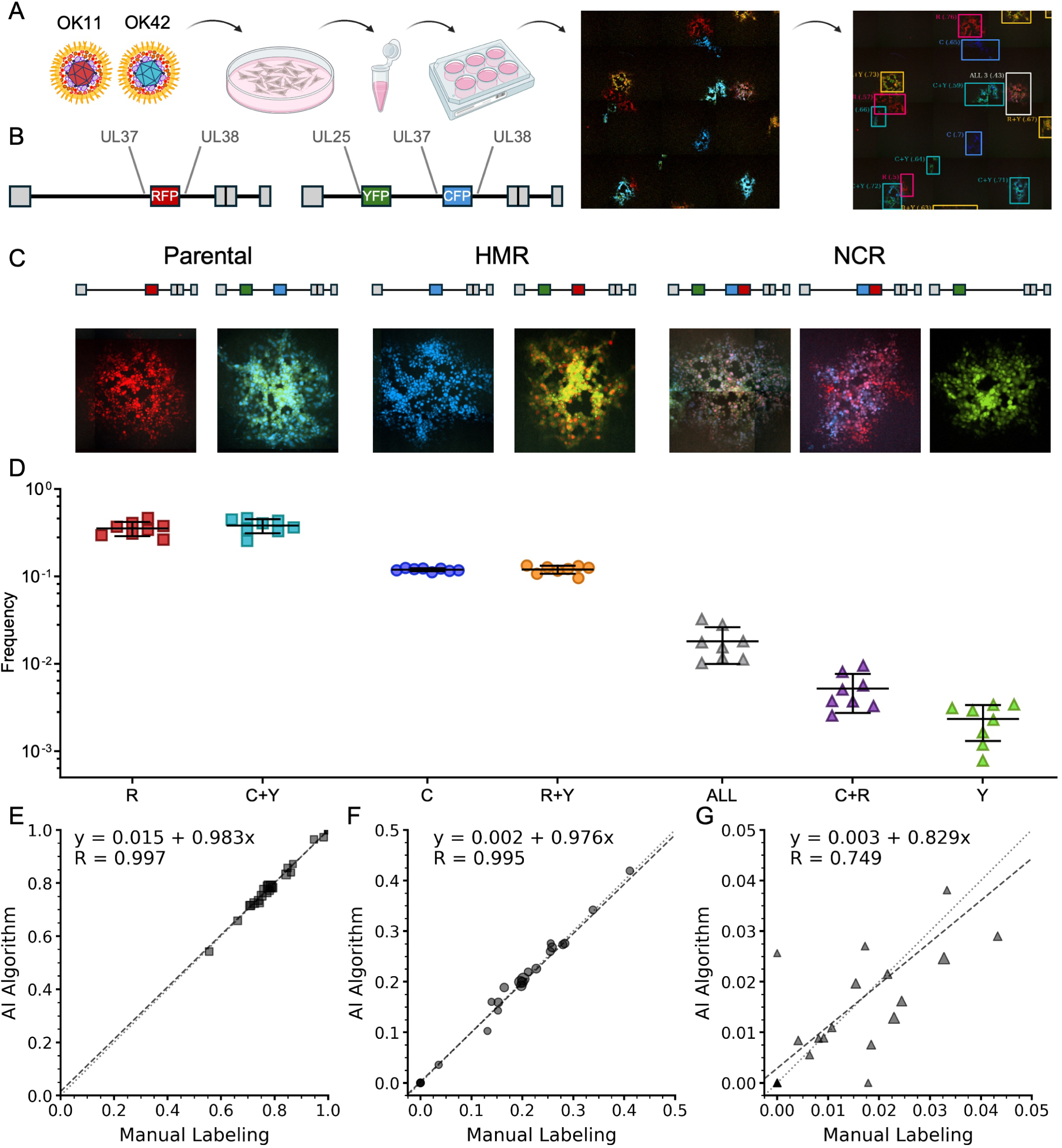
Experimental HSV-1 recombination assay. **A.** Schematic overview of three-color recombination quantification assay. HFF Cells are infected with an equal mix of HSV-1 expressing either mCherry (RFP; OK11) or mTurq2 and EYFP (CFP and YFP; OK42). 18 HPI released progeny are collected and replated on Vero cells for individual plaques. Plaques are then imaged, classified and quantified using our deep learning-based object detection model (Plaquer). **B.** Schematic representation of OK11 and OK42 genome insertions. **C.** Schematic illustration of the seven possible viral progeny (parental, HMR and NCR) each with an imaged example plaque. **D.** Frequency of plaques in each class. Each dot is calculated from a single experiment (8 experiments, a total of 48,397 plaques were analyzed) and represents the number of plaques in a class to total plaques ratio. Horizontal line represents the mean, and error bars indicate standard deviation. **E-G.** Object detection and classification model compared to manual labeling performance. Each point represents calculated rates of one large image. Shape size is relative to the total number of plaques in the image. A total of 23 large images were evaluated. Performance in **E.** parental, **F.** HMR and **G.** NCR is described by linear regression analysis, with fitting equations and Pearson’s correlation coefficients (R). Dotted lines have a slope of 1 while dashed lines display the linear fit.

To quantify recombination events, we scanned thousands of progeny plaques produced by coinfection at different conditions. We developed a deep learning model to detect, classify and count plaques according to their colors (see Methods and **Figure 1A,C and D**). 255 large images were manually annotated and counted by several researchers. 232 of these images (47,577 plaques in total) were used for the development process and 23 images were set aside for final model evaluation.

Parental, HMR and NCR rates were calculated as the ratio of number of plaques in each group, relative to the total number of plaques. Performance was evaluated by comparing manually analyzed with model predicted rates of Parental, HMR and NCR of each image. Model predicted rates of Parental and HMR groups were highly correlated with the manual rates (R=0.997 and R=0.995 for Parental and HMR, respectively; **Figure 1E and F**). For the NCR group, the model was less in agreement with the manual counts (R=0.749, **Figure 1G**). This may be attributed to the rarity of NCR events and subsequent susceptibility to variation resulting from a single plaque labeling difference.

### Recombination rates assay detects both HMR and NCR differences

To verify that our recombination assay can identify differences in rates of HMR and NCR events, we tested the effect of multiplicity of infection (MOI) on recombination rates. It was previously shown that intergenomic HMR rates are highly dependent on MOI (31). We coinfected immortalized human foreskin fibroblasts (HFF) cells with an equal mix of HSV-1 OK11 and HSV-1 OK42 at different MOIs. 18 hours post infection (HPI), progeny viruses were collected, plated on Vero cells for individual plaques and analyzed for recombination rates. Significantly higher rates of HMR were observed as MOI increased, from 11% at MOI 0.2 to about 25% at MOI 20 (**Figure 2A**). NCR rates also increased with MOI, but this trend was not statistically significant. This is probably due to the rare nature of NCR events, only about 1.5% of progeny viruses at MOI 20 (**Figure 2B**), thus NCR rates are calculated based on few plaques per each image. These results suggest that our system, although noisy for NCR, can detect fractions of percent differences in recombination rates.

**Figure 2.**
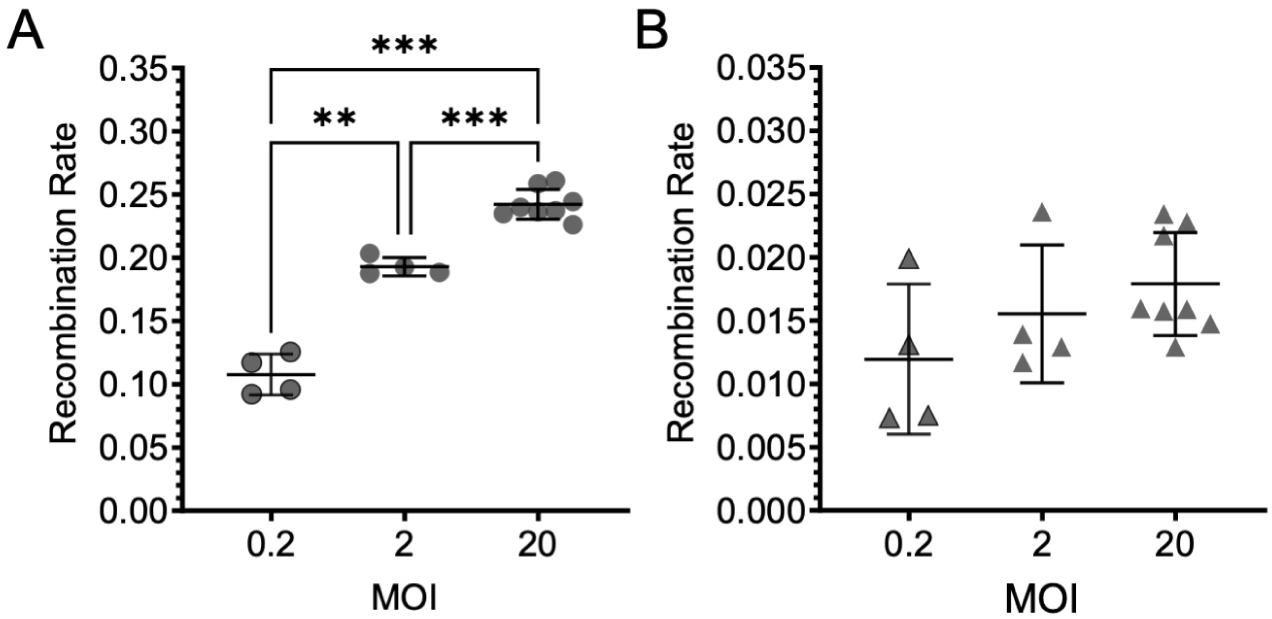
Viral recombination rate is MOI dependent. **A.** HMR rate of at MOI 0.2, 2 and 20. **B.** NCR Rate at MOI 0.2, 2 and 20. Data represent at least two independent experiments with at least two replicates each. Statistical significance was determined by Mixed Effects Analysis (*\*\*<0.01* and *\*\*\*<0.001*). Horizontal line represents the mean, and error bars indicate standard deviation.

### Limited effect of inhibitors of host proteins on viral recombination rates

Several host proteins were found to be associated with the HSV-1 replication forks. As some of these proteins are likely to be involved in viral recombination, we tested a handful of specific inhibitors of proteins known to be associated with the viral replication.

The host DNA clamp, proliferating cell nuclear antigen (PCNA), is a processivity factor involved in host DNA replication and recruitment of host DNA damage and repair factors (32). PCNA was identified to be associated with HSV-1 replication forks (25). PCNA-I1, a PCNA inhibitor, has been recently shown to reduce HSV-1 titers during infection and increase the production of defective virions (33). We therefore wished to measure the effect of PCNA inhibition by PCNA-I1 on recombination rates. At the three different concentrations we tested, the viral titers were reduced by 1 to 2.5 log compared to the untreated infection (**Figure 3A**). At 0.25 µM of PCNA-I1, although we observed a log reduction in titer, we detected a significant increase of about 10% in viral HMR rates (**Figure 3B**). At the higher concentrations in which the titer was reduced by 2 logs or more, HMR rates were similar to the untreated cells. No effect of PCNA-I1 was observed on NCR rates (**Figure 3C**).

**Figure 3.**
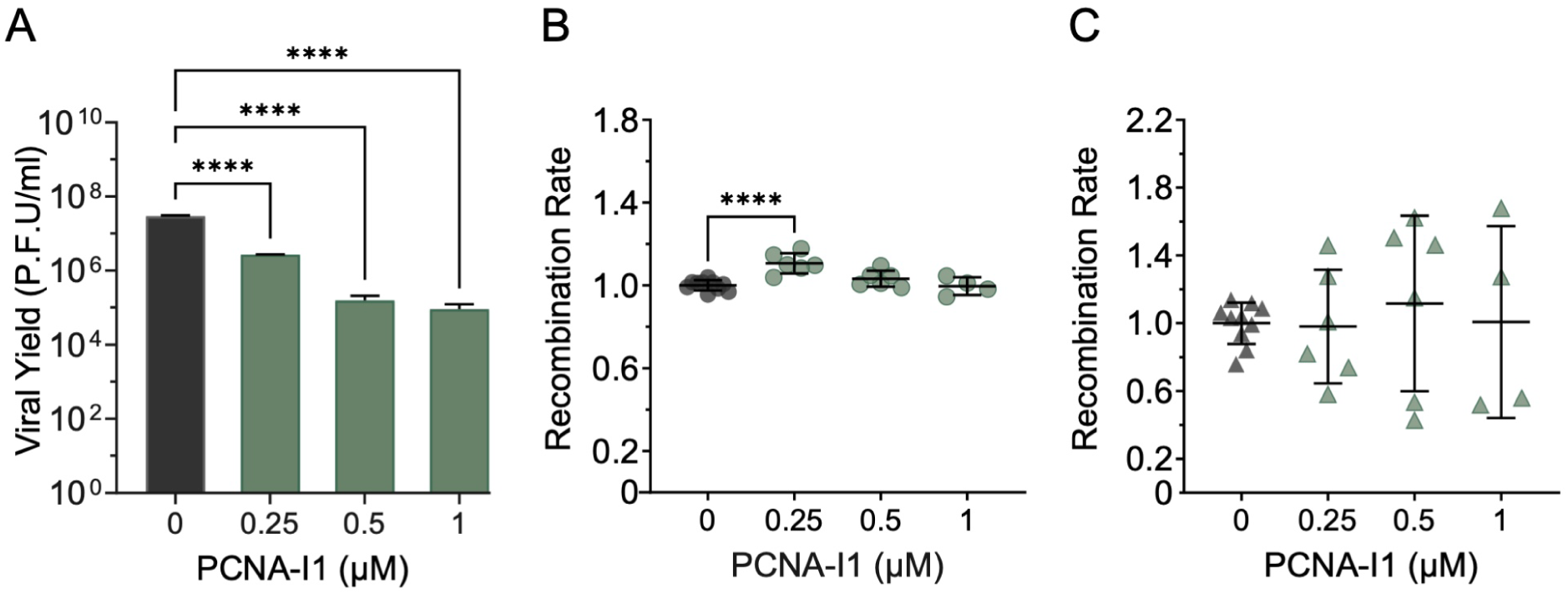
PCNA inhibitor reduces viral yield and partially increases the rate of homology mediated recombination. HFF cells were infected at MOI 20 in the presence of PCNA-I1. **A.** Viral yield in the presence of PCNA-I1. Bars represent the titer mean from one experiment with two biological replicates and four technical replicates each. **B-C.** Normalized **B.** HMR and **C.** NCR rates in the presence of PCNA-I1. Each circle represents recombination rates normalized to corresponding experiment control mean. Data represent three independent experiments with at least two biological replicates each. Statistical significance was determined by Ordinary One-way ANOVA (****<0.0001). Horizontal line represents the mean, and error bars indicate standard deviation.

We selected inhibitors for two proteins known to be associated with viral replication forks: PARP-1 and Mre11, and two proteins with transient association with the replication forks as they are substrates of the viral ICP0 protein: TP53BP1 and DNA-PK (24–26, 34–36). All these proteins are associated with the host response to DNA damage, mainly to double-strand DNA breaks (37). We carried out the coinfection recombination assay on HFF cells at MOI 20 as described above in the presence of each inhibitor (See Table 1 for the list of inhibitors) at 3 different concentrations. None of these inhibitors had shown a consistent dose dependent effect on viral titers or viral recombination rates **(Figure S1**).

**Table 1.** List of inhibitors used in this work.

| Target | Inhibitor | Source |
| --- | --- | --- |
| Mre11 (MRN complex) | Mirin (52) | Merck |
| PARP-1 | 3-Aminobenzamide (3AB)<br>(53) | Sigma |
| TP53BP1 | UNC9512 (54) | MedChemExpress |
| DNA-PK | NU7441 (55) | Tocris |
| PCNA | PCNA-I1 (33) | Sigma |
| Viral polymerase | Acyclovir (ACV) | Sigma |
| Viral polymerase | Phosonoacetic acid (PAA) | Sigma |
| Viral helicase | Amenamevir | Cayman Chemical |
| Viral helicase | Pritelivir | Sigma |
| Cellular ribonucleotide reductase | Hydroxyurea | Shiloh Lab, Tel Aviv University, Tel Aviv. |

### Viral replication inhibitors increase homology mediated recombination rates

Although HMR is generally regarded as tightly linked to viral replication, our observations with PCNA-I1 suggest that partial inhibition of replication can lead to elevated HMR rates. To test the effect of viral replication inhibition on the HMR rates, we tested several viral inhibitors that target the replication complex. We tested two viral polymerase inhibitors: Acyclovir (ACV) and Phosphonoacetic acid (PAA), and two helicase-primase inhibitors: Pritelivir (PRT) and Amenamevir (AMN). All four inhibitors at three different concentrations were tested for their effect on the rate of recombination following coinfection of HFF cells at MOI 20. All HMR rates were significantly higher in the presence of the different inhibitors at all concentrations; increased by 24 to 29% for ACV, 25 to 31% for PAA, 13 to 24% for PRT and 18 to 29% for AMN (**Figure 4A-D**). In contrast, almost all NCR rates were significantly reduced in these conditions by 43 to 58% for ACV, 57 to 61% for PAA, 41% (only at the highest concentration) for PRT and 40 to 42% for AMN (**Figure S2A-D**). However, results of the NCR were not as consistent as the HMR results (not in all concentrations) and at the low concentration of PRT we even detected a 28% increase in NCR rate. At these concentrations of drugs, viral titers were reduced by between 85 to 99.2% for ACV, between 78 to 99.4% for PAA, between 34 to 90% for PRT and between 82 to 99.1% reduction for AMN (**Figure 4E**). These results suggest that viral replication inhibitors can increase recombination rates under conditions that reduce progeny yield by up to two logs, whereas higher concentrations limited progeny recovery and prevented meaningful analysis.

**Figure 4.**
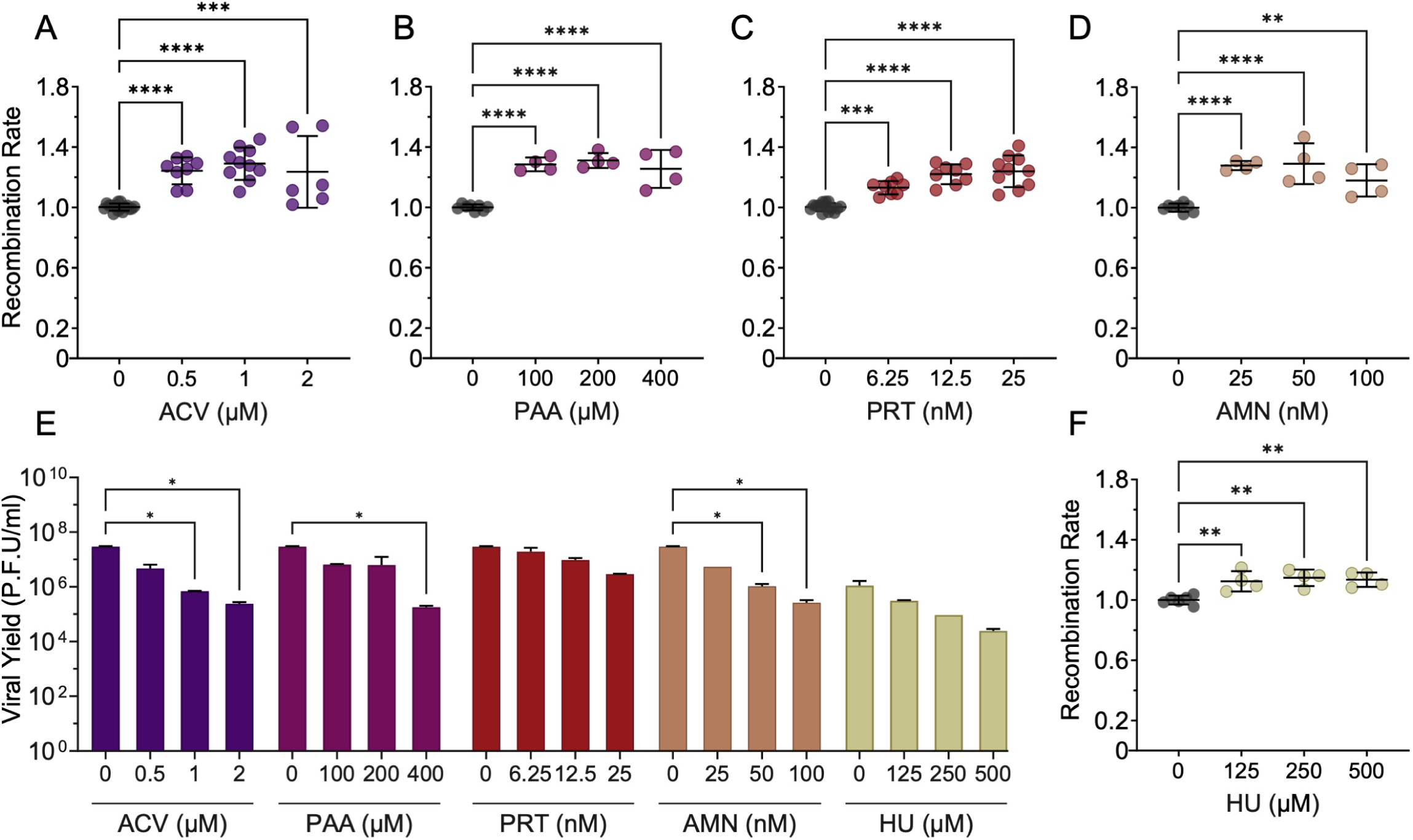
Inhibitors of viral replication increase the rate of viral homology mediated recombination. HFF cells were infected at MOI 20 in the presence of inhibitors of viral replication. **A-D, F.** Normalized HMR rates for specific inhibitors of viral DNA replication: **A.** Acyclovir, **B.** Phosphonoacetic acid, **C.** Pritelivir, **D.** Amenamevir and **F.** Hydroxyurea. Each circle represents HMR rates normalized to corresponding experiment control mean. Data represent at least two independent experiments with at least two replicates each. Statistical significance was determined by ordinary one-way ANOVA (*\*\*<0.01, ***<0.001* and *\*\*\*\*<0.001*). Horizontal line represents the mean, and error bars indicate standard deviation. **E.** Viral yield in the presence of indicated viral replication inhibitors. Bars represent the titer mean from one experiment with two biological replicates and four technical replicates each. Error bars indicate standard deviation. Statistical significance was determined by ordinary one-way ANOVA (\**< 0.05*).

To test whether the increase in viral HMR is unique to inhibition of viral replication proteins, we also tested the effect of hydroxyurea (HU), which reduces deoxyribonucleotides pools in the cells, indirectly inhibiting viral DNA replication. At the three concentrations tested, HU increased the HMR rates by 12 to 15% (**Figure 4F**), but had no effect on NCR rates (**Figure S2E**). At these concentrations, viral titer was reduced by 72 to 98% (**Figure 4E**). These results suggest that moderate inhibition of viral replication process by any mechanism increases the rates of HMR.

To verify that this increase in HMR is not cell type specific, we repeated the experiments with ACV and PRT on human immortalized retinal pigment epithelial (RPE-1) cells. RPE-1 cells were infected at MOI 20 with the two viral constructs in the presence or absence of the drugs. At 18 HPI, progeny viruses were collected and plated on Vero cells for plaque formation, as described previously. Analysis of plaque fluorescence indicated that both ACV (27–35%) and PRT (16–26%) increased HMR rates at all concentrations tested (**Figure 5A-B**). Unlike HFF cells, in the RPE-1 experiments the NCR rates were not significantly decreased (**Figure S3A-B**), yet an increase (35%) in NCR rates was observed in the lower concentration of PRT similar to the results in HFF cells. The drug concentrations used in the RPE-1 experiments reduced viral titers between 97.31% to 99.68% for ACV and 83.19% to 96.77% for PRT, resembling their effect on the viral yield in HFF cells (**Figure 5C**). These results indicate that partial inhibition of HSV-1 replication (less than 3 log reduction in titer) increases HMR rates both in fibroblasts and in epithelial cells.

**Figure 5.**
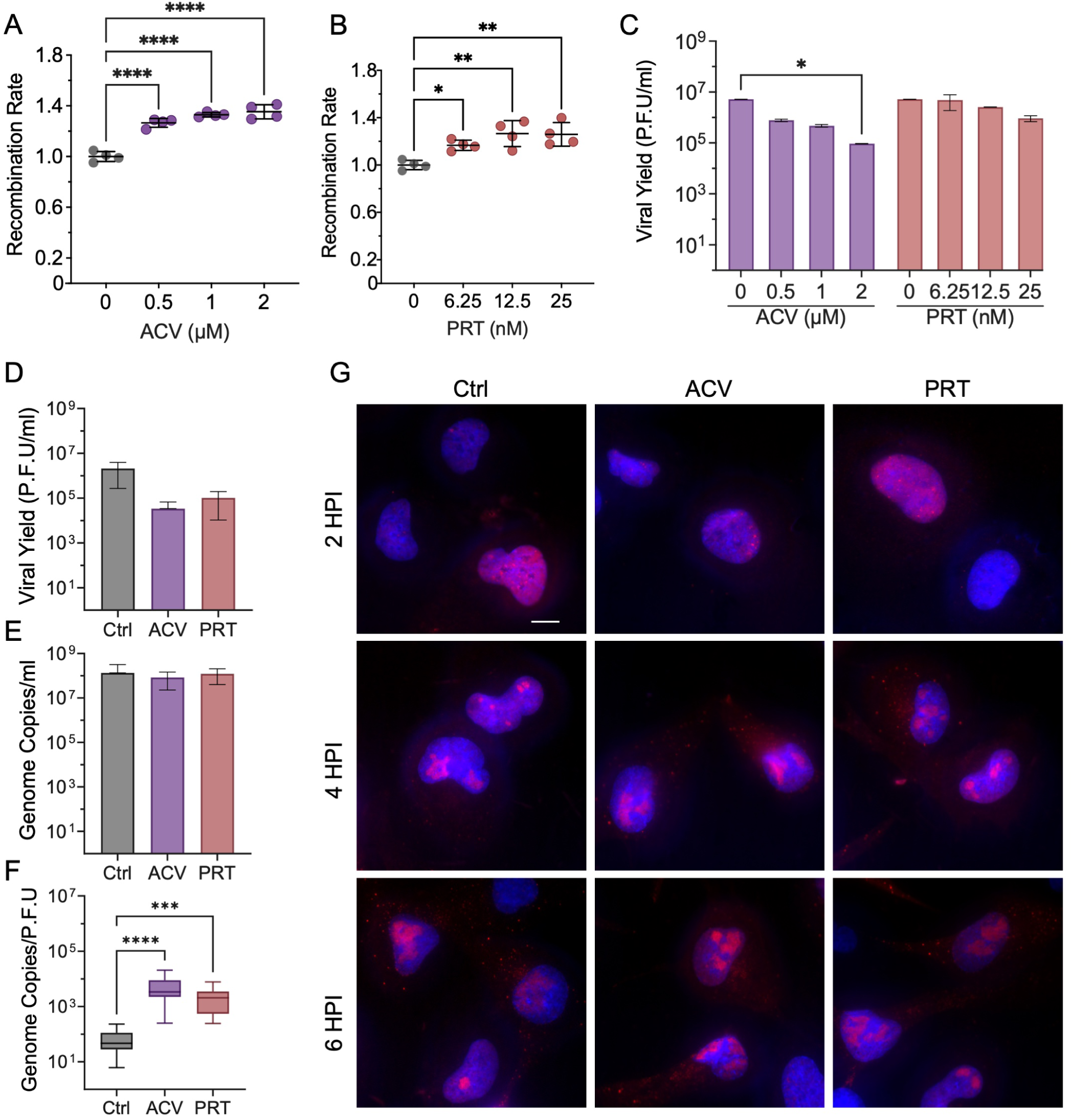
Inhibitors of viral replication reduce viral yield without affecting rate of replication. RPE-1 cells were infected at MOI 20 in the presence of inhibitors of viral replication. **A-B** Normalized HMR rates for **A.** Acyclovir and **B.** Pritelivir respectively. Each circle represents HMR rates normalized to corresponding experiment control mean. Data represent two independent experiments on RPE-1 cells with two replicates each. Statistical significance was determined by ordinary one-way ANOVA (*\*\*<0.01, ***<0.001* and *\*\*\*\*<0.001*). Horizontal line represents the mean, and error bars indicate standard deviation. **C.** Viral yield from RPE-1 cells in the presence of Acyclovir and Pritelivir, respectively. Bars represent the titer mean from one experiment with two biological replicates and four technical replicates each. Error bars indicate standard deviation. Statistical significance was determined by ordinary one-way ANOVA (\**< 0.05*). **D-F.** RPE-1 cells infected with HSV-1 strain 17+ in the presence of 2 µM Acyclovir or 25 nM Pritelivir. Bars represent the mean from two experiments with two biological replicates and four technical replicates each. Error bars indicate standard deviation. **D.** Viral yield, **E.** DNA copy number and **F.** Genome-to-PFU ratio were measured. Statistical significance was determined by ordinary one-way ANOVA for D and E, Kruskal-Wallis ANOVA for F (*\*\*\*<0.001* and *\*\*\*\*<0.001*). **G.** RPE-1 cells infected with OK42 in the presence of 2 µM Acyclovir or 25 nM Pritelivir, fixed at 2, 4 and 6 HPI and stained for ICP4 (red). Nucleus stained with DAPI (blue). Scale bar = 10 µm.

### Partial inhibition of HSV-1 replication decreases viral infectivity

To test whether increased recombination events resulted in more defective viral genomes, we measured the genome-to-PFU ratio in the presence of ACV and PRT. RPE-1 cells were infected with HSV-1 strain 17 at MOI 20 in the presence or absence of either 2 µM ACV or 25 nM PRT. 18 HPI the viruses were collected, titered and analyzed by qPCR for viral genomes. Similar to the recombinant analysis, drug treatment reduced viral titers by 98.6% with ACV and 95.1% with PRT (**Figure 5D**). In contrast, DNA copy numbers decreased by only 54.9% with ACV and 34.2% with PRT (**Figure 5E**), raising the genome-to-PFU ratio from ∼45 in untreated samples to ∼1200 with PRT and ∼3200 with ACV (**Figure 5F**). These results indicate that at these drug concentrations, reduced viral titers reflect the accumulation of defective genomes or particles rather than impaired genome replication.

To further evaluate the effect of the viral inhibitors on viral replication, we visualized viral replication compartments within the RPE-1 cells at different time points post infection. Infection was carried out using HSV-1 strain 17 in the absence or presence of either 2 µM ACV or 25 nM PRT. Viral replication compartments were visualized by immunofluorescence using an anti-ICP4 antibody. The antiviral treatment did not result in any significant differences in the size or distribution of the replication compartments represented by the ICP4 transcription activator puncta (**Figure 5G**). From these findings, we suggest that antivirals targeting the herpes replication machinery at moderate concentrations reduce viral progeny primarily by impairing infectivity, with only a limited effect on genome replication.

### Viral replication inhibitors increase interspecies recombination

HSV-1 and HSV-2 are closely related viruses with similar genomic organization. Recent evidence has demonstrated their ability to recombine in vivo (7, 8, 38). We speculate that the mechanism of interspecies recombination resembles that of intraspecies recombination and is therefore likely to be similarly influenced by the antiviral drugs tested above. To test this hypothesis, we coinfected HFF cells with HSV-1 ND2 (expressing GFP-ICP4 and mRFP-VP26) and the HSV-2 G strain. Cells were infected at MOI 20 with a mix of the two viruses in the absence or presence of 2 µM ACV or 25 nM PRT, and progeny were collected at 18 HPI. The viruses were replated for plaque formation, with single-color plaques (green or red) considered recombinant (**Figure 6A**). The mean frequency of single-color plaques was 3.2% in untreated cells, increasing to 3.6% (14.5% increase) and 4.3% (37.2% increase) with ACV and PRT, respectively (**Figure 6B**). Thus, in agreement with our finding on intraspecies recombination, partial inhibition of the viral replication machinery increased interspecies recombination rates.

**Figure 6.**
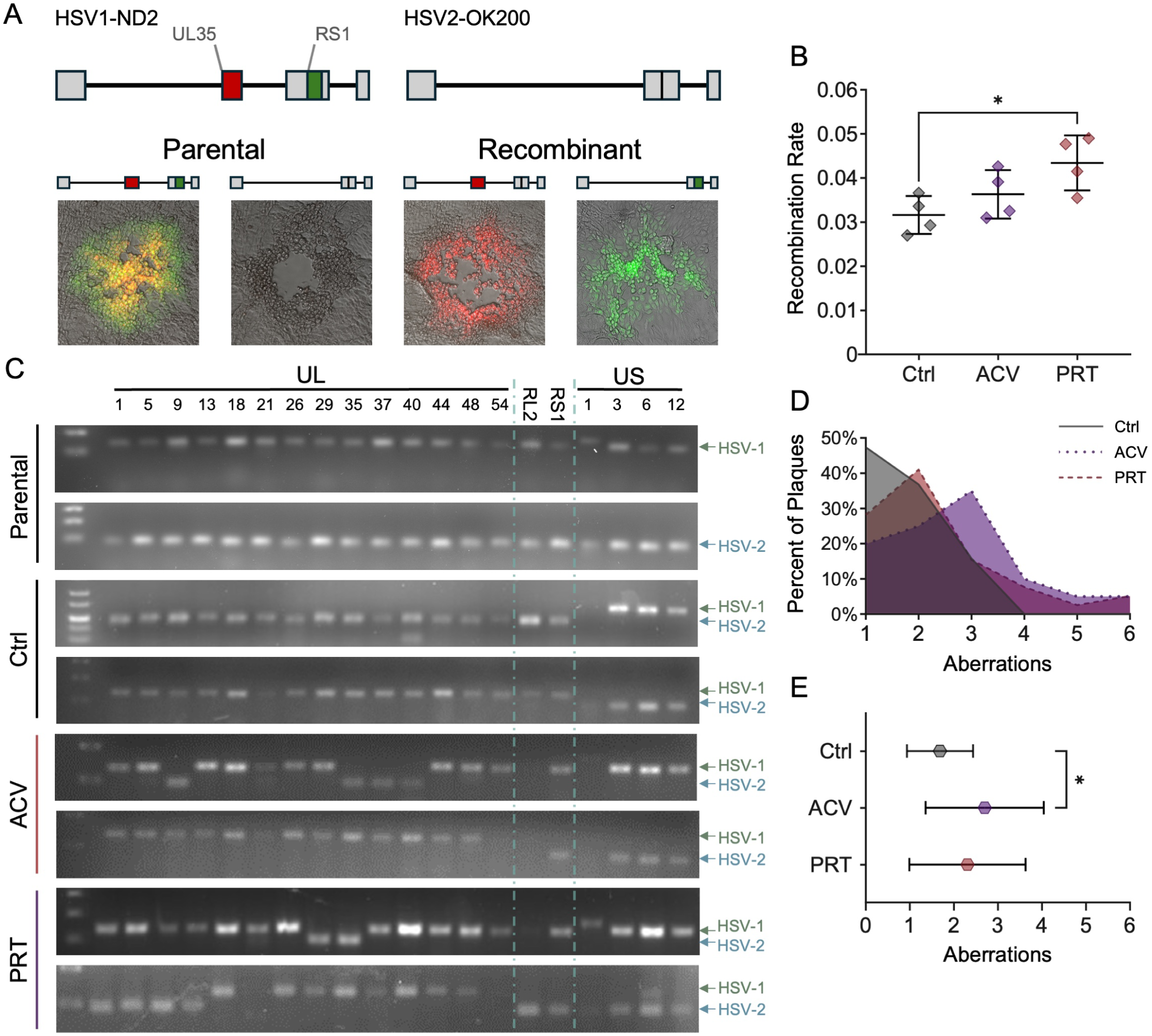
Viral polymerase inhibitors increase the rate of recombination between HSV1 and HSV2. **A.** Schematic representation of HSV-1 expressing ICP4-YFP and VP26-RFP (ND2) and HSV-2 strain G and possible viral progeny (parental, recombinants) each with an example plaque. **B.** Recombination rates in the presence of 2 µM Acyclovir (purple) or 25 nM Pritelivir (red) compared to control (gray) on HFF cells. Recombination rates were calculated by manually classifying dual-, single- and no-FP plaques. Data represent two independent experiments with two replicates each. Between 300 to 2735 plaques were classified for each replicate. Statistical significance was determined by ordinary one-way ANOVA (\**<0.05*). Horizontal line represents the mean, and error bars indicate standard deviation. **C.** Single-FP plaques from B were purified and analyzed from two independent experiments with two replicates each. Example PCR analysis for 2 purified recombinant single-FP plaques from either control (out of 19), Acyclovir (out of 21) or Pritelivir (out of 39) initial experiment. Expected specific amplicons for HSV-1 (green arrow) and HSV-2 (blue arrow) was ∼135bp and ∼94bp respectively. Example PCR for HSV-1 and HSV-2 was performed on lysed viral ND2 and OK200 stocks, respectively (note that HSV-1 US1 has a larger PCR fragment than expected). Dashed turquois line annotates the repeat genes; RL2 and RS1. **D-E.** For each plaque, the number of observed aberrations (crosses, deletions and duplications) detected by the PCR assay (**C**) was summed, **D.** Frequency distribution of aberrations per plaque and **E.** the average number of aberrations per plaque, were plotted for each treatment. Colored as in **B.** Statistical significance was determined by ordinary one-way ANOVA (\**<0.05*).

We exploited the use of two different viruses to test whether viral replication inhibitors increase genetic aberrations in progeny viral genomes. To do so, we first plaque purified 19, 20 and 39 single-color plaques from the drug free, ACV treated and PRT treated coinfections, respectively. The purified recombinant viruses were then analyzed by 20 specific primer sets for either HSV-1 or HSV-2, distributed every 5,000 to 10,000 bp along the genomes (14 primer sets within the U_L_ segment, 4 primer sets in the U_S_ segment, and 1 primer set for each repeat region, as specified in **Tables S1 and S2**). As shown in **Figure 6C**, HSV-1 PCR products are larger than those of HSV-2, enabling us to distinguish between the two genomes at each site tested. For our purified plaques, we detected three different types of aberrations: recombination crosses, deletions and gene duplications (a positive PCR for both HSV-1 and HSV-2). Most deletions and duplications were verified twice to prevent random PCR failure. All three types of aberrations were observed with or without the drugs (**Figure 6C**). To quantify aberrations per genome, we used a conserved approach that considered the minimal number of events necessary to account for our findings. This included counting the fewest possible crosses across all four genomic isomers, combining deletions when feasible, excluding duplications in repeat-region PCR products due to the presence of two repeat copies, and disregarding crosses within repeats that may simply represent correction with the corresponding repeat during plaque purification. We observed up to three aberrations per plaque in control infections and up to six aberrations under both drug treatments (**Figure 6D**). The mean number of aberrations per plaque was 1.68 in untreated samples, increasing to 2.7 with ACV (P = 0.0125) and 2.31 with PRT (P = 0.0512) (Figure 6E). Taken together, these findings suggest that antiviral drugs targeting viral replication increase both the frequency of interspecies recombination and the number of genomic aberrations, consistent with our earlier observations in intraspecies events.

## Discussion

The role of viral recombination as a major driver of genetic diversity in herpesviruses populations has become increasingly evident with the advent of next-generation sequencing (NGS) technologies (10, 39). However, identifying specific factors that contribute to viral recombination independently of replication has remained a challenge, as these processes are tightly intertwined (19, 21, 40). Here, we developed a fluorescence-based recombination assay coupled with a deep learning model for image analysis, enabling quantification of viral intergenomic homology-mediated and non-canonical recombination rates (HMR and NCR, respectively). We found that inhibitors of host proteins that we tested had almost no effect on recombination rates, whereas antiviral drugs that inhibit viral replication increase the rates of HMR significantly. We observed a similar effect of the antiviral drugs on interspecies recombination (between HSV-1 and HSV-2).

A key advantage of our assay is its capacity to analyze hundreds of thousands (more than 450,000 plaques were analyzed) of progeny plaques, enabled by Plaquer, our deep learning-based object detection model. Deep learning (DL), a subset of machine learning, employs neural networks to learn complex data representations (see Methods). Unlike conventional image processing algorithms that rely on predefined parameters, such as object size, fluorescence intensity, and absolute thresholds values, DL models learn directly from specific data, enabling robust generalization to previously unseen examples (41). Similar to other DL models that were shown to outperform human experts in consistency and agreement across various tasks, our model yielded highly reproducible results, enabling reliable quantification of HMR rates across large sample sizes and diverse experimental conditions (**Figure 1E-G**).

We have recently demonstrated that HSV-1 NCR can be detected both in culture and in vivo (11). Given the prevalence of NCR events, we designed our assay to estimate their frequency alongside HMR. Our assay can detect only very specific NCR events, either gene duplication or deletions of the fluorescent proteins, that likely represent only a small fraction of all NCR events. Due to their markedly lower frequency compared to HMR events (**Figure 1**), NCR rates were often noisy, resulting in a difficulty in detecting significant, reproducible differences under most conditions (**Figure 2 and S1-S3**). Previously, we suggested that gene duplications are unstable, as plaques carrying them could not be purified (17). This hypothesis, that duplications occur frequently but represent a transient state, is consistent with the genomic accordion model proposed for vaccinia virus (42).

Many host proteins interact with the HSV-1 replication forks (25, 26). Several of these proteins (including: PCNA, Mre11, PARP-1, and to lesser extent TP53BP1) are not recruited to the viral replication forks in the absence of the viral exonuclease UL12 (suggested to be involved in viral recombination) (43). However, in our experiments most host-targeting inhibitors did not consistently alter viral recombination or replication (**Figure 3 and S1**). These results may be attributed to redundancy in protein function, which could limit the impact of inhibiting any single factor. An alternative explanation is that the inhibitors do not block the specific functions required by HSV-1, as observed with the PCNA inhibitor T2AA (33).

In our experiments, the only host protein inhibitor that reduced viral titers was PCNA-I1 (**Figure 3E**). PCNA-I1 is known to stabilize the PCNA homotrimer, impairing its ability to load or unload from DNA. Packard et al. have shown that PCNA-I1 does not inhibit PNCA loading to viral DNA yet it reduced viral replication compartment size and increased the DNA to PFU ratio (33). To ensure sufficient progeny for recombination analysis, we used lower concentrations of PCNA-I1 than those reported by Packard et al. Despite this, all concentrations tested resulted in a significant reduction in viral titer. Notably, only the lowest concentration led to a significant increase in HMR rates (**Figure 3F**), mirroring the effect observed with viral replication inhibitors (**Figure 4**). We speculate that higher concentrations of PCNA-I1 reduces replication to a degree that masks measurable changes in HMR rates. A similar phenomenon was observed with high concentrations of ACV, PAA and AMN, where viral titers were reduced by approximately two logs, and recombination rates became more variable, with smaller increases in HMR rates compared to lower drug concentrations (**Figure 4A,B,D,E**). These results suggest that, in our system, significant increases in HMR rates are detectable only under conditions of moderate replication inhibition.

Viral polymerase inhibitors (ACV and PAA) and viral helicase-primase inhibitors (PRT and AMN) increased HMR rates at concentrations that reduced viral titers by up to two logs in two different cell lines (**Figures 4 and 5A-C**). At these concentrations, our data suggest that DNA replication is only minimally affected, as indicated by comparable replication compartment sizes and genome copy numbers between treated and untreated infections (**Figure 5E,G**). We propose that partial inhibition of replication, i.e. replication stress, can be alleviated through recombination of stalled genomes (44). This is further supported by our observation that secondary replication inhibition induced by HU treatment, which depletes the nucleotide pool, also led to an increase in HMR (**Figure 4F**). However, not all treatments that reduce viral titers are likely to increase recombination rates. For example, Tang et al. (2014) reported that siRNA-mediated knockdown of cellular recombination factors Rad51 and Rad52 led to reduced HSV-1 titers and decreased recombination rates (28).

Interspecies recombination between HSV-1 and HSV-2 is increased in the presence of ACV and PRT (**Figure 6B**), suggesting a mechanism similar to that driving intraspecies HMR, although baseline interspecies recombination rates are substantially lower. Moreover, the antiviral drugs increased other aberrations in the progeny viral genomes (**Figure 6C-E**) consistent with the elevated genome-to-PFU ratio observed for HSV-1 (**Figure 5F**). These results suggest that the genetic rearrangements occurring during infection are more complex than observed by specific genetic markers and by analyzing only the plaque-forming viral genome.

Enhancing HMR may offer practical advantages for both research and therapeutic applications. Many recombinant herpesviruses used in experimental systems are generated through HMR (45), and increasing its efficiency could improve the production of such constructs. Recent work has also proposed the use of gene drive– based HSV recombinants to target latent viral genomes (46). The increase in HMR rates observed with replication-inhibiting antivirals suggests that these small-molecule inhibitors could potentially accelerate gene drive propagation.

ACV is frequently used to establish latency in vitro (47), typically at concentrations much higher than those employed here and intended to completely block replication. Nevertheless, our results suggest that the effects of ACV on latent genomes and progeny viral genomes should be further investigated in such models.

The concentrations of 2 μM (∼0.45 mg/L) ACV and 25 nM (∼10 μg/L) PRT used in this study fall within the lower range of reported therapeutic serum levels in humans (48). Importantly, effective concentrations in infected tissues are likely to be even lower. Additionally, since ACV is (in many countries) available over the counter in topical formulations, its use may not always adhere to recommended clinical guidelines.

These factors imply a potential clinical relevance of the concentrations examined in our experiments.

These concentrations of ACV were used recently to follow the accumulation of drug resistance genomes during in vitro passaging (48). Yet, the likelihood that drug-resistant viral genomes would emerge and alter the outcome of our experiments (single 18-hour infections in the presence of the drug) is extremely low, as even after five passages at this concentration, the population IC50 did not increase significantly (48).

In recent years, an increasing number of HSV-1 and HSV-2 intraspecies recombinants have been detected in clinical settings (38). While this rise is most likely attributable to improved detection methods and greater anatomical overlap of infections, our findings suggest that antiviral treatment may also contribute to recombination and viral diversification.

Our findings should not be interpreted as questioning the safety or clinical value of these antiviral therapies. Rather, they suggest a potential unintended consequence of reduced drug concentrations that warrants further investigation, particularly regarding their impact on viral evolution and the risk of drug resistance.

## Methods

### Cells

Vero, African green monkey kidney epithelial cells [ATCC CCL-81; American Type Culture Collection (ATCC), Manassas, VA, USA] and HFF, h-TERT immortalized human foreskin fibroblasts (a kind gift from Sara Selig Technion, Haifa) were grown with Dulbecco’s Modified Eagle Medium (DMEM; biowest), supplemented with 10% Fetal Bovine Serum (FBS; Gibco) and 1% penicillin (10,000 units/ml) and Streptomycin (10 mg/ml) (Pen-Strep; biowest). RPE-1, immortalized Retinal Pigmented epithelial cells (a kind gift from Eran Bacharach Tel Aviv University, Tel Aviv) were grown with F12 (Gibco) supplemented with 10% FBS, 1% Pen-Strep and 0.02% Hygromycin B (50mg/ml, Toku-E). All cells were grown at 37°C with 5% CO2.

### Viruses

All viral constructs expressing fluorescent proteins are derivatives of HSV-1 strain 17+. HSV-1 OK11 carries a single fluorescent protein (mCherry) with a nuclear localization tag under the CMV promoter between UL37 and UL38 genes (29). HSV-1 OK42 was constructed by coinfection of previously described HSV-1 MT01 (carries a single fluorescent protein (mTurq2) with a nuclear localization tag under the CMV promoter between UL37 and UL38 genes (49)) and HSV-1 OK19 constructed with EYFP gene fused in-frame within the UL25 gene after the 50th amino acid of the viral protein (50). ND02 was kindly provided by Nir Drayman (The University of California, Irvine) and carries YFP-ICP4 and mRFP-VP26 fusions (51). HSV-2 strain G [VR-3393] was obtained from ATCC.

### Recombination Assay

12-well plates were seeded with HFF or RPE-1 cells. Two hours pre-infection, medium with or without inhibitors (see **Table 1** for details) at the indicated concentration was added. Cells were inoculated with an equal mix of OK11 and OK42 at MOI 20 (unless indicated otherwise) and incubated at 37°C for 1 hour. Following incubation, inoculum was removed, washed three times with PBS and fresh medium with or without inhibitors was added. After incubation for 18 hours at 37°C, only the medium was collected and stored at -80°C. Samples were sonicated and centrifuged to pellet any cell debris and the supernatant was diluted to infect confluent Vero cells in 6-well plates at semisolidified medium.

### Imaging

36-48 hours post infection 4 areas were imaged from each well. Viral Plaques and nuclei were visualized using Nikon Eclipse Ti-E epifluorescence inverted microscope (Nikon, Tokyo, Japan).

### Plaquer

Viral plaques in 255 three-color images (resolution ranging between 5300^2^ and 6144^2^ pixels) were manually annotated using Nikon NIS Elements AR (v4.2.0). Images were split into train, validation and test in a ∼70-20-10 ratio, with plaque class stratification. Two models were developed for plaque detection and quantification, each designed to optimally detect plaques of different sizes. Each image was divided into an 8×8 grid with 50% overlap (225 sub-images) for the first model, and into a 4×4 grid with 50% overlap (49 sub-images) for the second model. To tackle the class imbalance in the data, data augmentations were implemented. For each small patch, affine transformations were performed, and synthetic patches were randomly generated with rare plaques. Each model was based on fine-tuning YOLOv8 (Ultralytics). Following the prediction for each small patch, a final prediction was required for the large images from both models. Non-max suppression (NMS) was used for stitching the patch-level predictions. Predicted bounding boxes contained by others were discarded, as well as abnormal predictions based on bounding boxes’ aspect ratio and area. For each image, the model outputs the total count of plaques in each class and the original image with the predicted bounding boxes (**Figure 1A**).

### DNA copies to PFU

RPE-1 cells were incubated with Acyclovir or Pritelivir at 2 µM and 25 nM, respectively, 2 hours pre infection. Cells were inoculated with HSV-1 strain 17+ at MOI 20 and incubated for 1 hour at 37°C. Inoculum was removed and medium with or without inhibitors was added. At 18 HPI, medium was collected. Samples were sonicated and simultaneously prepared for qPCR and plaque assay to quantify PFU/ml. For qPCR, each sample was incubated in lysis buffer (10 mM Tris-HCL pH=7.5, 1 mM EDTA, 150 mM NaCl, 1% Triton-X100; Sigma, 0.04% Proteinase K; NEB) at a 1:1 ratio for 30 minutes at 37°C then 10 minutes at 95°C for Proteinase K inactivation. For PFU/ml standard plaque assay was performed on Vero cells.

### Quantitative PCR (qPCR)

For viral DNA standard curve pure HSV-17+ was serially diluted in Ultra-Pure Water (UPW; Gibco) starting at 3.8 ng/µl and all lysed samples were diluted 1:100 in UPW. Three sets of primers were used (HSV-1 UL13, UL44 and US3, see Primer table) in a reaction mix containing Fast AP Sybr Green mix (Applied Biosystems), 0.5 µM of each primer and UPW. 1 µl diluted DNA sample was added per reaction. Samples were quantified using a CFX Duet Real-Time PCR System (Bio-Rad).

### **I**mmunofluorescence

Cover slips pre-treated with 0.1% poly-L-Lysin (Sigma) were seeded with RPE-1 cells. After pre incubation with or without the drugs, cells were inoculated with HSV-1 strain 17+ at MOI 20 at 37°C for 1 hour. Cells were washed, medium with either 2 µM Acyclovir, 25 nM Pritelivir or without drugs was added and cells were incubated at 37°C. At 2, 4 and 6 HPI, cells were washed with PBS, fixed with 4% Paraformaldehyde (PFA; bioworld) for 15 minutes, then washed again with PBS. Cells were permeabilized for 10 minutes with 0.5% Triton-X, washed with PBST (0.01% Triton-X in PBS) and blocked for 30 minutes with 5% Bovine Serum Albumin (BSA; WVR) in PBST. Slides were incubated 2 hours with HSV-1 ICP4 primary antibody (HSV-1 ICP4 (10F1) sc-56986; Santa Cruz). Slides were washed with PBST and incubated with CFL555 secondary antibody (anti-mouse CFL555 sc-362267; Santa Cruz) for 1 hour. Slides were mounted with Mounting media with DAPI (Applied Biosystems), sealed and visualized using Nikon Eclipse Ti-E epifluorescence inverted microscope (Nikon, Tokyo, Japan).

### HSV-1/2 Recombination Assay

HFF cells were infected with an equal mix of viral constructs ND2 and HSV-2 strain G in the presence or absence of 2 µM ACV or 25 nM PRT as described above. Progeny viruses were collected at 18 HPI, and plaques were imaged and manually counted according to the color.

Single-colored plaques (RFP or YFP) were picked and purified for two or three rounds. A lysate was prepared from each purified plaque. For each plaque, PCR was performed with all sets of primers. Specific primers were designed for every 5 to 10Kbp for either HSV-1 or HSV-2 so that the expected PCR products were ∼135 bp or ∼94 bp, respectively (**Tables S1 and S2**).

### Statistical analysis

Graphs and statistical analysis were performed using GraphPad Prism 10.3. The statistical analysis of Plaquer performance was done with Python (version 3.11) and SciPy (version 1.13.1).

## Data Availability

All data is presented in the paper. Plaquer code can be found at GitHub and weights at Zenodo. Further information or clarification can be provided upon request from the corresponding author.

## Acknowledgement

We thank all Kobiler lab members for their comments. This work was supported by grant from the Israel Science foundation (grant # 1816/21) to O.K. The funders had no role in study design, data collection and analysis, decision to publish, or preparation of the manuscript.

